# Variability in mineral composition of Canadian lentil cultivars

**DOI:** 10.1101/2024.05.14.592802

**Authors:** Ana Vargas, Rajib Podder, Maya Subedi, Kirstin E. Bett, Albert Vandenberg

**Affiliations:** Dep. of Plant Sciences, Univ. of Saskatchewan, 51 Campus Dr., Saskatoon, SK S7N 5A8; Agriculture and Agri-food, Canada Saskatoon, SK, S7N 0X2

**Keywords:** lentils, mineral, trace element, chemical composition

## Abstract

Lentils are a good source of essential minerals for the proper functioning of the human body. We evaluated 34 cultivars and elite lentil lines representing the breadth of the Canadian breeding program. Trials were established in 10 site-years across Saskatchewan Province. Concentrations of 27 minerals were quantified with an inductive coupled argon plasma emission spectrometer in whole and dehulled lentil seeds. Li, V, Cr, Co, As, Ag, Cd, Sn, La, Hg, and Pb had concentrations below the quantification limit and were excluded from further analysis. The effects of site year, tissue type (whole and dehulled), and lentil genotypes were analyzed using a mixed model. Mineral concentrations of Na, K, Ca, Mg, Cu, B, and Ba were stable across site years. Na, Zn, P, Cu, Se, and Mo had similar concentrations in whole and dehulled seeds. Ca, Fe, Mg, Mn, B, Al, and Ba were more concentrated in whole seeds, while K, S, and Ni were higher in dehulled seeds. Several lentil genotypes had outstanding concentrations of several minerals. Lentil genotypes with a higher composition of several minerals could be a starting point for enhancing mineral composition in lentils.

## INTRODUCTION

Pulses are a crucial component of sustainable diets because of their nutritional profile and have well-characterized agri-environmental benefits. They have been consumed as a staple food in developing nations for thousands of years and are gaining attention in developed nations for their potential positive effect on human health (Olmedilla et al., 2010). The demand for plant-based protein will continue to grow, forecasting that these products will replace 20% of the meat and dairy consumed (National Research Council Canada 2023). Lentils (*Lens culinaris* Medik.) in particular, have been an essential component of the human diet for millennia. They are high in essential nutrients and require a short cooking time resulting in smaller nutrient losses than those with a harder seed coat (Satya et al., 2010). Lentils are a great source of protein – some varieties containing upwards of 30% (Wang and Daun 2006) and represent an attractive source for people choosing plant-based diets. Lentils are also a good nutritional option for people with restricted diets due to their low glycemic index (Campos de la Vega et al., 2010). Lentils are also an excellent source of minerals (Chen et al., 2021), and studies have described pulses as a potential whole food source for people suffering from micronutrient malnutrition (Thavarajah et al., 2009).

The human body requires various minerals for its proper functioning. These requirements include bone and brain development, muscle building, and heart function. Lentils contain many minerals that are vital to the human body. Iron (Fe) is essential for most living organisms for oxygen and, electron transport, and DNA synthesis (Lieu et al., 2001). Calcium (Ca) is the most prevalent mineral in the human body, and its metabolism includes other nutrients such as vitamin D, phosphorus (P), and protein (Beto 2015). Zinc (Zn) is the third most abundant mineral in the body and has an extensive function in wound repair and reproduction (Maxfield and Crane 2020). Phosphorus is an essential part of cellular structures and is important for maintaining cellular functions (Razzaque 2011). Potassium (K) plays an important role in maintaining cellular function and is the most abundant cation in intracellular fluid (Stone et al., 2016). Selenium (Se) is required for the proper functioning of the immune system (Rayman 2000). Magnesium (Mg) is an essential cofactor in over 300 enzymatic reactions (Schwalfenberg and Genuis 2017), and regulates muscle contraction, blood pressure, and is necessary for the synthesis of DNA, RNA, and proteins (Grober et al., 2015).

Half of the human population has dietary deficiencies of essential mineral elements which leads to cardiovascular disease and imbalance in many biological pathways (White and Broadley 2009). Fe deficiency causes many health conditions including anemia, inability to maintain body temperature, and mortality of pregnant women (Boccio et al., 2003). Zinc deficiency is characterized by delayed growth and impaired immune function (Prasad 2004). Copper deficiency is responsible for neurological problems like neuropathy (Scheiber et al., 2013), while Mg deficiency can lead to abnormal heart rhythms, and epileptic seizures among other conditions (Yuen et al., 2012). Manganese deficiency can contribute to birth defects, impaired infertility, and skeletal malformations (Aschner et al., 2002), while K deficiency is associated with cardiac arrest and weakness (Soar et al., 2022).

Over the last two decades, efforts have been made to quantify mineral contents in lentils, specifically Fe, Zn, Ca, K, P, Mg, Mn, Cu, and Se. Many of these studies include Canadian lentil genotypes grown in different parts of the world (Chen et al., 2022; Vandemark, et al., 2018; Gupta et al., 2016; Ray et al., 2014; Wang et al., 2011; Thavarajah et al., 2009; Wang and Daun 2006), as well as Mediterranean genotypes (Plaza et al., 2021) and middle Eastern lentil material (Haq et al., 2011; Hefnawy et al., 2011).

With increasing interest in pulse consumption, obtaining information on individual mineral concentration and establishing the mineral profile of the newest Canadian lentil materials is warranted. Industry partners have also expressed interest in mineral profiles. Sodium (Na), potassium (K), and calcium (Ca) are of particular interest to food and ingredient companies because they serve as flavor-potentiating ions and are usually included in the nutrient facts panel as part of food labeling requirements.

Lentils are also a good source of other less studied minerals which are, thus far unquantified in lentil cultivars. An inductively coupled argon-plasma emission spectrometer (iCAP 6500; Thermo Jarrell Ash Corp., Franklin, MA, USA) was used to assess 27 minerals and trace elements in Canadian cultivars and elite lines from the lentil breeding program of the Crop Development Centre (CDC) at the University of Saskatchewan. Our objectives were to establish a full profile of the mineral composition of recent lentil cultivars from the CDC, to compare these to older cultivars, and to determine the best breeding approach moving forward for enhancing the mineral composition in lentils.

## MATERIALS AND METHODS

### Plant material and field conditions

Thirty-four cultivated genotypes (cultivars and lines) were grown in 5 locations during 2017 and 2018 for a total of 10 site-years across Saskatchewan. The locations were Sutherland 17 (52°09’58.2” N, 106°30’21.8” W), Rosthern 17 (52°41’21.1” N, 106°18’00.6” W), Elrose 17 (51°17’59.0” N, 107°59’29.0” W), Lucky Lake 17 (51°03’58.3” N 107°11’30.6” W), Limerick 17 (49°38’28.12“N, 106°29’15.91“W), Sutherland 18 (52°09’57.7“N, 106°30’15.7“W), Rosthern 18 (52°41’16.71“N, 106°17’57.20“W), Elrose 18 (51°18’3.06“N, 107°59’5.37“W), Lucky Lake 18 (51° 3’57.94“N, 107°11’34.74“W) and Limerick 18 (49°38’28.12“N, 106°29’15.91“W). Trials were grown during the typical growing season-from May to August- in plots of 3 m^2^, three rows per plot, with inter-rowing spacing of 30 cm, a row length of 3 m, and a target population of 540 plants per plot. The experiments were arranged in a Randomized Complete Block Design (RCBD) with three replications. Experiments were conducted under rainfed conditions, and weeds and pests were controlled using the recommended local practices (Table 1). Genotypes and classification of their respective market classes are described in Table 1.

**Table 1.**
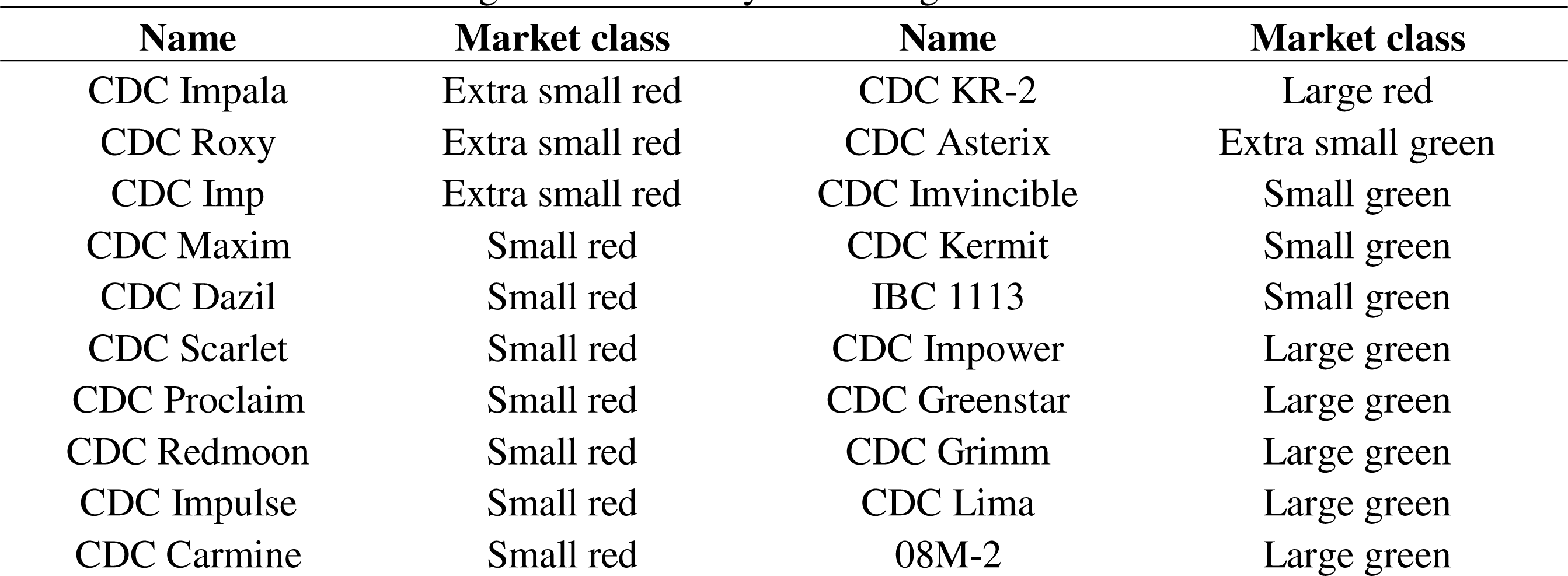
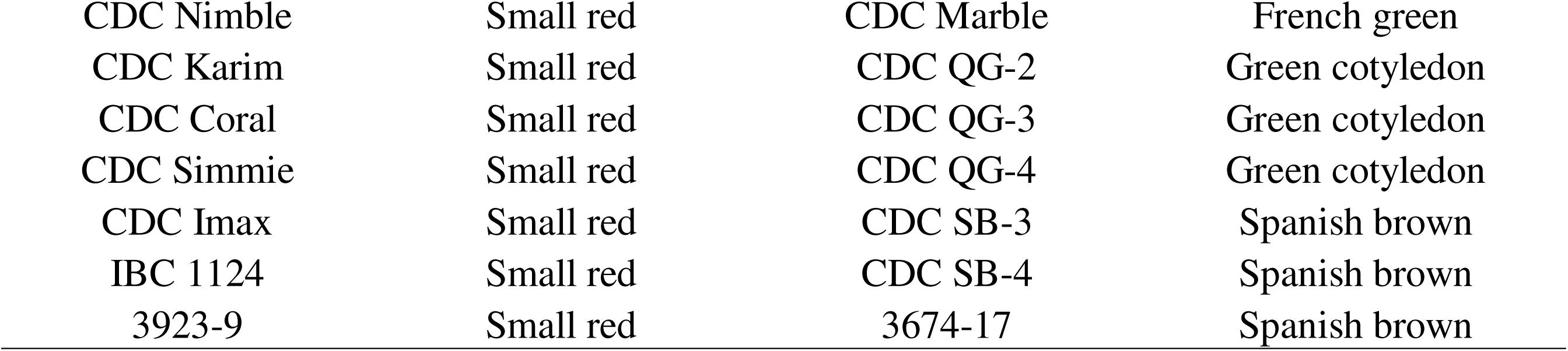
Name and market class of 34 lentil cultivars and breeding lines studied for their concentration of 27 minerals grown at 10 site-years during 2017-18 in Saskatchewan.

### Assessment of 27 mineral concentration in whole and dehulled lentils

Mineral analysis was conducted at the Crop Development Centre of the University of Saskatchewan, SK, Canada. Whole seeds and dehulled samples were evaluated to compare mineral concentration differences among both types, as these are the two forms lentils consumed. A Satake TM05C Grain Testing Mill (Satake Engineering Co. Ltd, Hiroshima, Japan) was used to dehull the whole lentils and after separating the hull, clean dehulled samples were used for the analysis. All samples were ground using a Udy grinder (Cyclone Sample Mill, Seedburo Co., Chicago, IL, USA).

Concentration of minerals was quantified with an inductively coupled argon-plasma emission spectrometer (iCAP 6500; Thermo Jarrell Ash Corp., Franklin, MA, USA) following the procedure of Glahn et al. (2017). Mineral concentrations of iron (Fe), zinc (Zn), sodium (Na), potassium (K), calcium (Ca), magnesium (Mg), phosphorus (P), sulfur (S), manganese (Mn), copper (Cu), selenium (Se), molybdenum (Mo), nickel (Ni), boron (B), aluminum (Al), and barium (Ba) were analyzed. Other minerals including lithium (Li), vanadium (V), chromium (Cr), cobalt (Co), arsenic (As), silver (Ag), cadmium (Cd), tin (Sn), lanthan (La), mercury (Hg), lead (Pb) was tested for, but their concentrations were below the limit of quantification, therefore, excluded from the analysis.

### Experimental Design and Statistical Analysis

A Levene’s test was conducted to determine the homogeneity of variance among site years. It was insignificant; therefore, data were treated as homogeneous and analyzed using a mixed model ANOVA in R (R Core Team 2021). Comparisons were made among lentil genotypes, tissue types, and site-years using a mixed model. The two tissue types analyzed (whole and dehulled seeds), and the 34 lentil genotypes were considered as fixed effects. Site-years were considered as one effect for a total of 10 site-years. Site-years and repetitions within site-years were considered as random effects. Means were separated with an LSD (least significant difference) test (P≤0.05). For differences among genotypes, results averaged across site years are presented.

A principal component analysis (PCA) was conducted to evaluate the effects of the site-years and tissue type to the variance observed across genotypes, and to explore any potential differences based on market class. Values were standardized before the PCA for equal contribution of each mineral to the analysis. The two principal components explaining the largest variation were plotted. A Pearson’s correlation analysis was conducted to determine if there were relationships between seed yield and mineral composition.

## RESULTS

### Mineral concentrations across different growing location

Significant differences in mineral concentrations among lentils grown in different locations for both whole and dehulled seeds (Table S2 for ANOVA). When the mineral concentrations across site-years were compared, Na, K, Ca, Mg, Cu, B, and Ba exhibited similar values for both whole and dehulled seeds (Figure 1, Table S3 for eigenvalues and correlations with original variables). Higher concentrations of Fe were observed at Sutherland 17, Sutherland 18, and Rosthern 18 for whole seeds. Similarly, higher concentrations of P and S were observed in whole seeds from Rosthern 17. Lentils from Elrose 17 and Elrose 18 had the highest concentrations of Mo (Figure 1a), but in contrast, had lower values for P and Mn. When the mineral concentrations for dehulled seed were compared, lentils grown at Rosthern 17 had the highest values across locations (Figure 1b). Lower concentrations of Zn were observed in whole and dehulled seeds in lentils grown at Elrose 17 (Figure 1).

**Figure 1.**
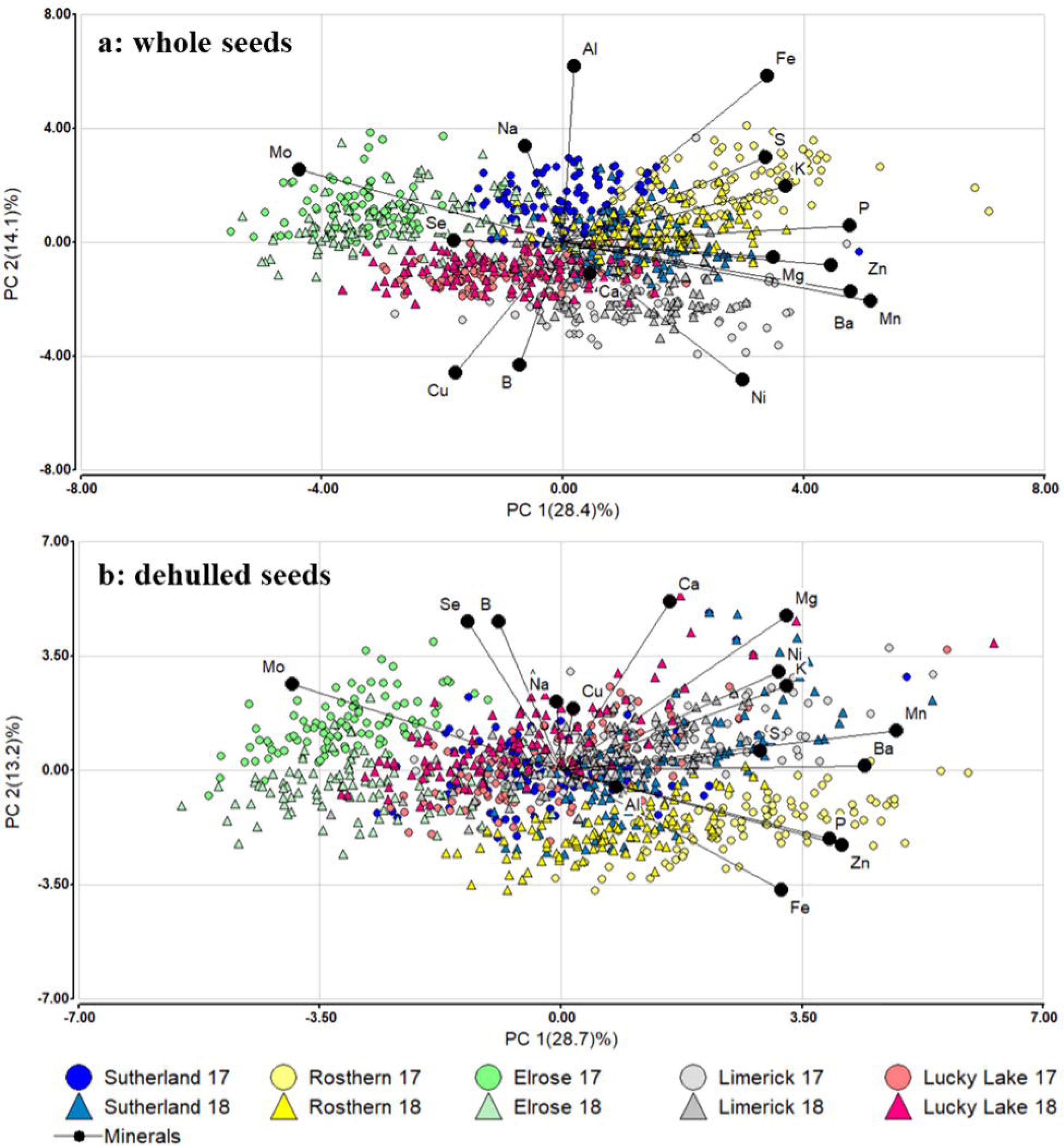
Principal component analysis of a: whole and b: dehulled seeds of 34 lentil genotypes according to 10 site-years grown during 2017-18 in Saskatchewan.

### Mineral concentrations in whole and dehulled seeds

The average mineral concentrations across site-years for whole and dehulled seeds were significantly different for some minerals (Table 2 for ANOVA). Na, Zn, P, Cu, Se and Mo had similar concentrations in both tissue types. Ca, Fe, Mg, Mn, B, Al and Ba had higher concentrations in whole seeds, while K, Ni and S had higher concentrations in dehulled seeds (Figure 2, Table S3 for eigenvalues and correlations with original variables).

**Figure 2.**
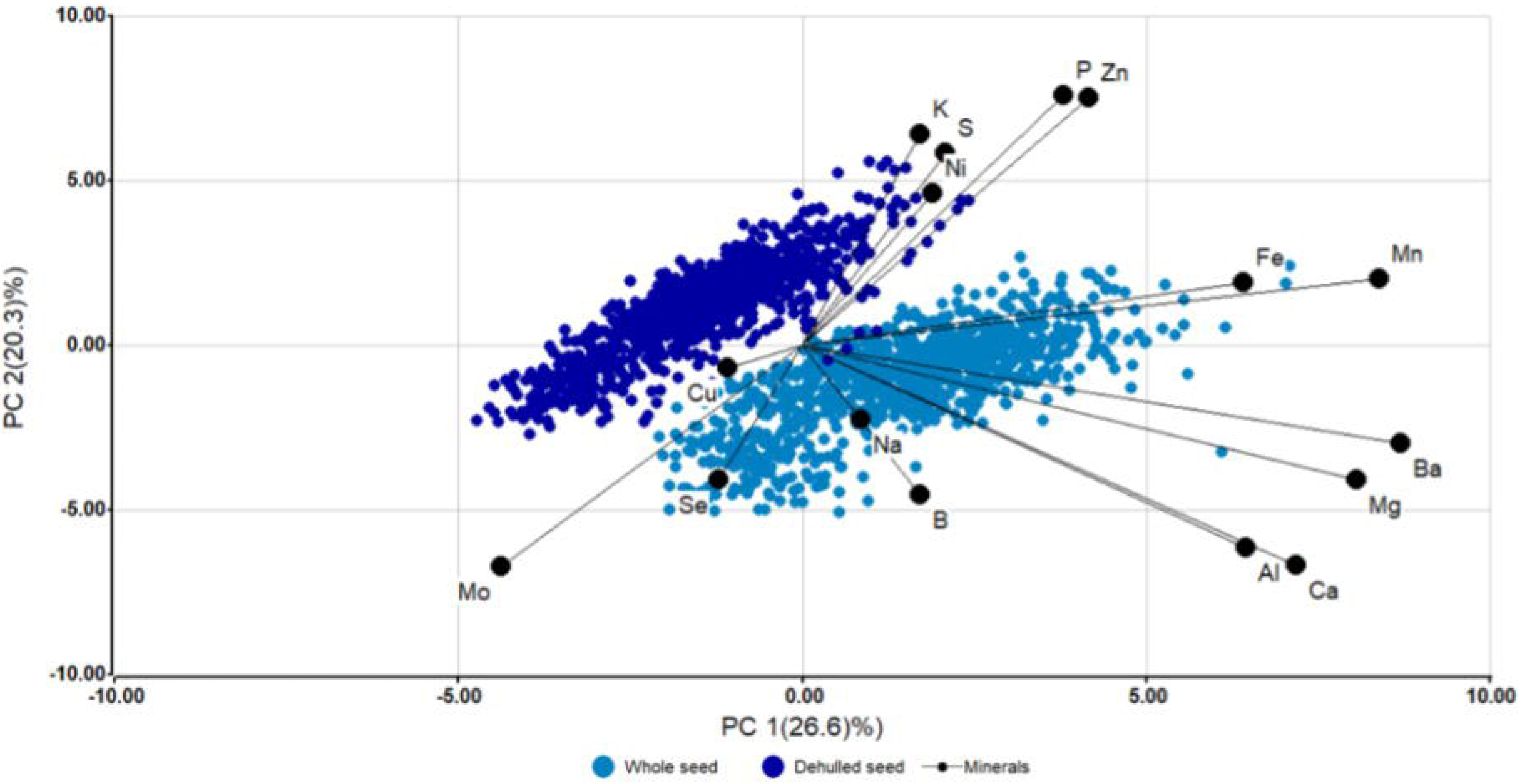
Principal component analysis of 34 lentil genotypes for whole and dehulled seeds grown in 10 site-years during 2017-18 in Saskatchewan.

**Table 2.** Average and range values for 16 minerals evaluated in whole and dehulled seeds of 34 lentil genotypes across 10 site-years in Saskatchewan during 2017-2018.

| Mineral | Average |  | Range |  | LSD |
| --- | --- | --- | --- | --- | --- |
|  | WS<sup>a</sup> | DS<sup>b</sup> | WS | DS |  |
|  |  |  | mg Kg<sup>-1</sup> |  |  |
| **Fe** | 64.8 | 57.0 | 51.6-72.5 | 43.1-64.4 | 0.41*** |
| **Zn** | 35.50 | 36.2 | 31.3-38.8 | 29.4-40.3 | 0.35*** |
| **Mg** | 991 | 749 | 932.5-1066.4 | 669.2-880.2 | 3.57*** |
| **Na** | 4.12 | 3.70 | 3.3-5.3 | 2.7-4.8 | 0.15*** |
| **Ca** | 643.4 | 250.0 | 472.5-821.7 | 171.8-328.3 | 3.87*** |
| **Mn** | 13.91 | 11.52 | 12.2-15.9 | 10.2-13.2 | 0.12*** |
| **Cu** | 7.89 | 7.90 | 7.2-9.1 | 6.9-9.0 | ns |
| **Mo** | 2.39 | 2.59 | 2.0-3.1 | 2.2-3.1 | 0.09*** |
| **B** | 7.65 | 6.55 | 6.6-8.7 | 5.6-9.2 | 0.13*** |
| **Ni** | 2.02 | 2.32 | 1.7-2.5 | 2.0-3.0 | 0.05*** |
| **Se** | 0.82 | 0.76 | 0.7-1.0 | 0.6-1.0 | 0.03*** |
| **Al** | 15.00 | 0.92 | 12.8-18.5 | 0.7-1.3 | 0.42*** |
| **Ba** | 5.30 | 1.72 | 4.1-6.4 | 1.4-2.2 | 0.07*** |
|  |  |  | g Kg<sup>-1</sup> |  |  |
| **K** | 8.95 | 9.29 | 8.4-9.5 | 8.6-9.9 | 0.03*** |
| **P** | 3.49 | 3.60 | 3.3-3.7 | 3.4-3.9 | 0.02*** |
| **S** | 1.89 | 1.95 | 1.7-2.0 | 1.8-2.1 | 0.01*** |

<sup>a</sup>WS: whole seeds, <sup>b</sup>DS: dehulled seeds. LSD: Least Significant Difference. ANOVA values are ns: not significant, \*, \*\*, \*\*\*: significant at 0.05, 0.01, 0.001 respectively.

We also looked at the range of mineral concentrations observed across genotypes (Table 2) for both whole and dehulled seeds. Ranges of variation were similar across genotypes in both tissue types. Between the lowest and the highest concentrations, Na, Ca, Ni and Mo had the broadest ranges of values, with a 1.5-fold increase from the lowest to highest genotypes. K and P had the least variation in concentration across genotypes while Fe, Zn, Mg, S, Mn, Cu, B, and Se concentrations ranged from 1.25-1.5-fold increase.

### Genotype specific mineral concentrations

Lentil genotypes varied in their mineral concentrations (Figure 3, Table S2 for ANOVA, Table S4 for all mean values) and those differences were generally independent of their market class (Fig. S1). Concentrations are reported in g Kg^-1^ for K, P and S, and in mg Kg^-1^ for all the other minerals.

**Figure 3.**
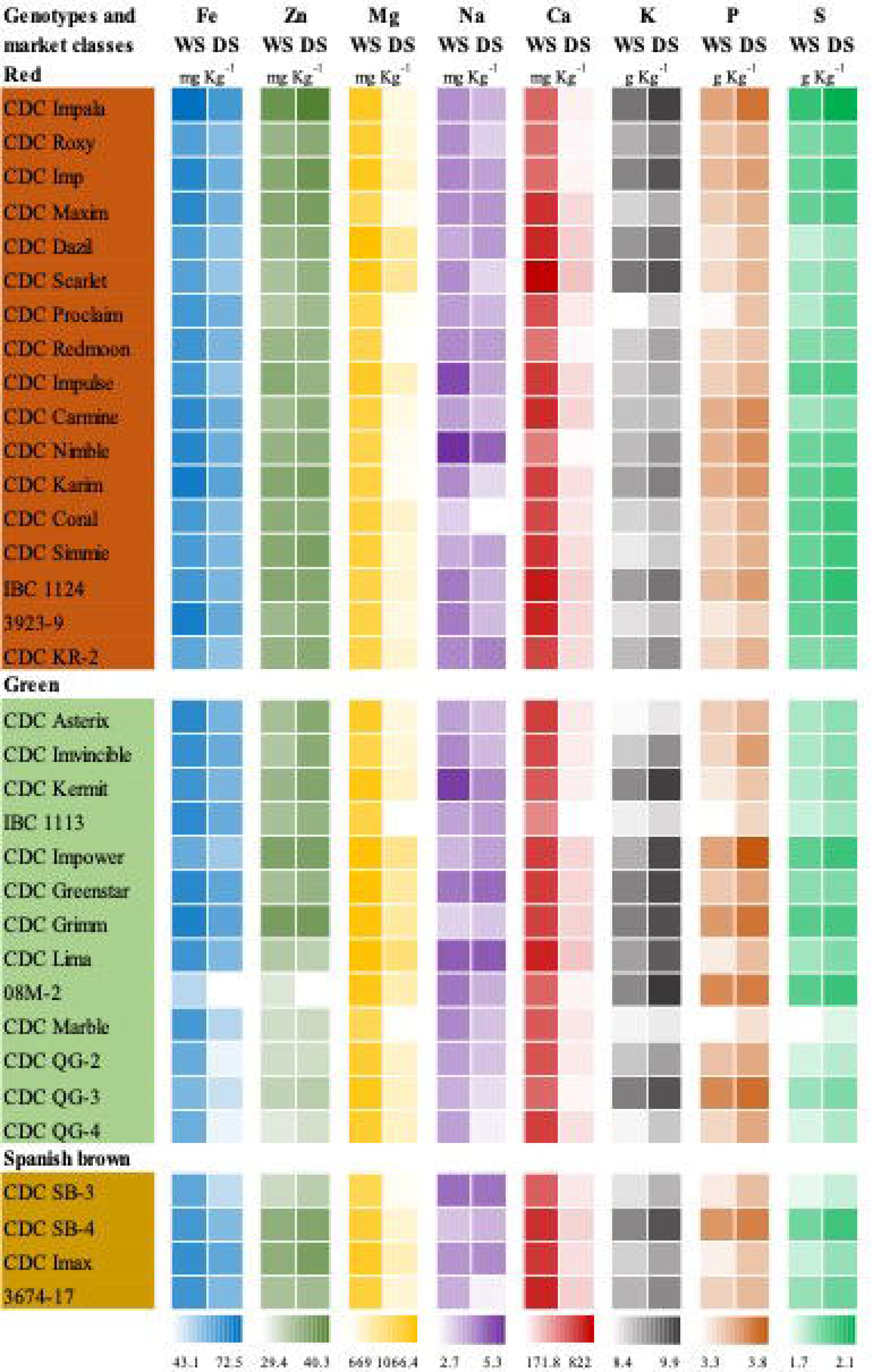
Concentration of 8 minerals in whole (WS) and dehulled seeds (DS) of 34 lentil genotypes evaluated across 10 site-years in Saskatchewan during 2017-18.

For Fe, ‘CDC Impala’ showed the highest concentration in whole seeds (72.5 mg Kg^-1^), compared to 08M-2, that had the lowest value in whole (51.6 mg Kg^-1^) and dehulled seeds (43.1 mg Kg^-1^). All green cotyledon cultivars ‘CDC QG-2’, ‘CDC-QG-3’ and ‘CDC QG-4’ had the lowest concentrations of Fe in dehulled seeds among all genotypes. Concentrations of Ca and Mg exhibited the greatest differences between whole and dehulled seeds, sometimes making the variation across genotypes greater between both tissue types. For instance, the Ca concentration in whole seeds of ‘CDC Scarlet’ was four times higher (821.7 mg Kg^-1^), compared to the concentration observed in the dehulled line IBC 1113 (171.8 mg Kg^-1^). Similarly, ‘CDC Dazil’ had twice the Mg concentration in whole seeds (1066.4 mg Kg^-1^), compared to the dehulled seeds of ‘CDC Marble’ (677 mg Kg^-1^) (Figure 3).

Some of the lentil varieties had significantly higher concentrations of Mn in whole seeds, including ‘CDC Karim’, ‘CDC Lima’ and ‘CDC QG-3’ (15.4, 15.9, 15.8 mg Kg^-1^ respectively), also higher than the average values observed across all genotypes when dehulled (11.5 mg Kg^-1^) (Figure 4, Table S4 for all mean values). Although the average concentration of B across genotypes was higher in whole seeds, some dehulled genotypes had similarly high concentrations. ‘CDC Proclaim’, ‘CDC Imvincible’ and ‘CDC Simmie’ showed the highest values (8.3, 8.7, 8.1 respectively) in whole seeds. Among dehulled seed samples, ‘CDC Impower’ and IBC 1124 had the highest concentrations of B (9.2 and 8.2 respectively) (Figure 4).

**Figure 4.**
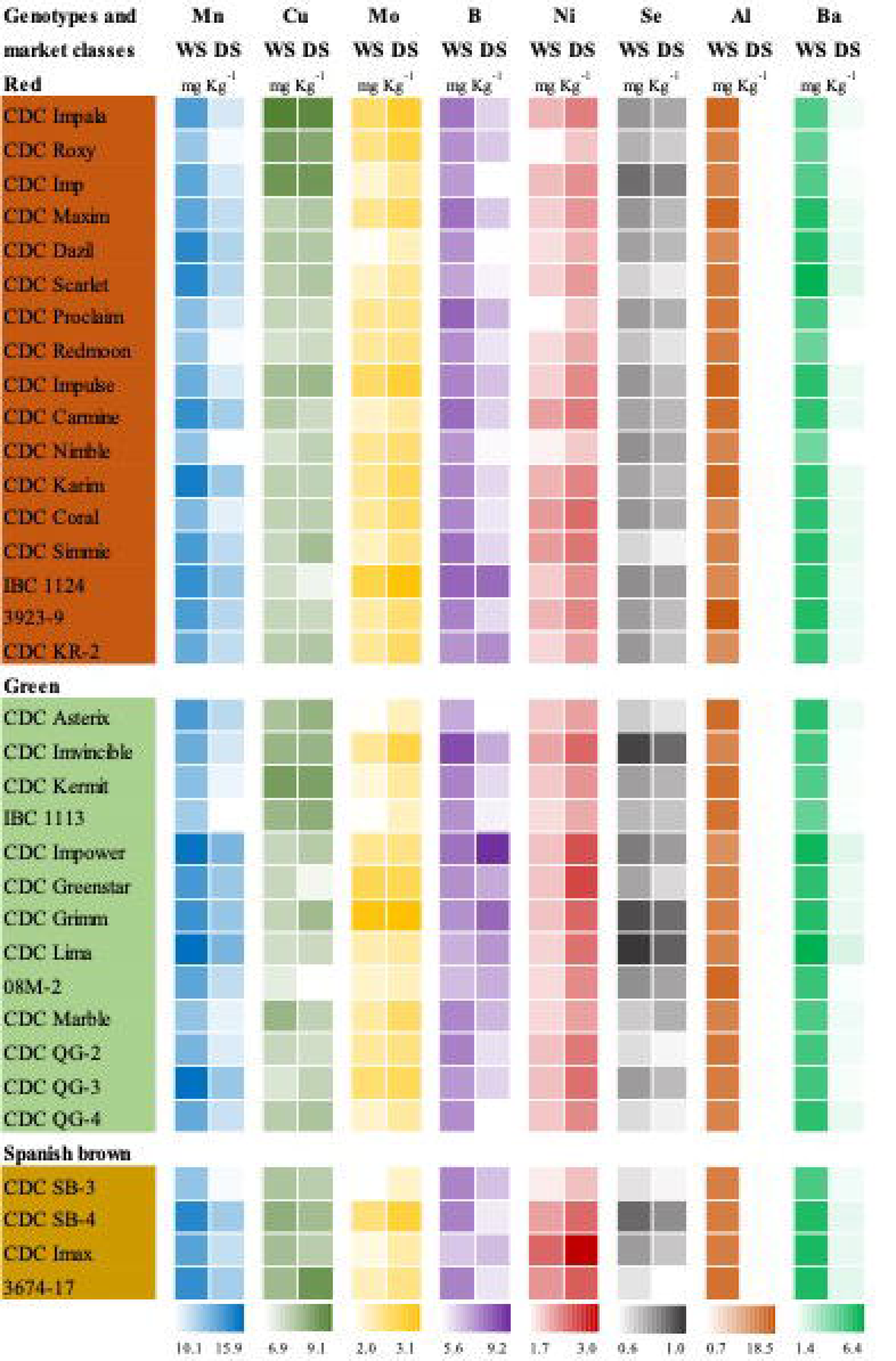
Concentration of 8 minerals in whole (WS) and dehulled seeds (DS) of 34 lentil genotypes evaluated across 10 site-years in Saskatchewan during 2017-18.

The S concentration in dehulled seeds of the cultivars ‘CDC Impala’, ‘CDC Impower’ and ‘CDC Grimm’ was the highest across genotypes in both tissue types (2.1, 2 and 2 mg Kg^-1^ respectively). In contrast, whole seeds of the cultivar ‘CDC SB-3’ had the lowest concentration of S (1.7 mg Kg^-1^) (Figure 3). Ni, which had also higher concentration in dehulled seeds, was the highest in seeds of the cultivars ‘CDC Greenstar’, ‘CDC Impower’ and ‘CDC Imax’ (2.6, 2.6, 3 respectively), contrasting with the values observed in whole seeds in ‘CDC Proclaim’ and ‘CDC Roxy’ (1.7, 1.8 respectively). This was also the case for the K concentrations observed, which were highest in the dehulled seeds of the cultivars ‘CDC Impala’, ‘CDC Asterix’ and 08M-2 (9.9 in all three) and significantly different from the K concentrations in whole seeds of the cultivars ‘CDC Proclaim’ and ‘CDC Asterix’ (8.4 in both) (Figure 4).

Among all the mineral concentrations, K was the most abundant element in the lentil samples. From the minerals that had similar concentrations in both tissue types, some specific genotypes exhibited higher values in whole seeds, dehulled seeds or both. Na concentrations were highest in whole seeds of ‘CDC Nimble’ and ‘CDC Kermit’ (5.3, 5.2 respectively) (Figure 3). Similarly, Se concentrations were highest in whole seeds of the cultivars ‘CDC Grimm’, ‘CDC Lima’, ‘CDC Imvincible’ and ‘CDC SB-4’ (1, 1, 1, and 0.9 respectively) (Figure 4).

The opposite was observed among the Zn and P concentrations where the highest values were in dehulled seeds (Figure 3). ‘CDC Impala’, ‘CDC Grimm’, ‘CDC Kermit’ and ‘CDC Asterix’ (40.3, 38.2, 37.3 and 37 respectively) has the highest concentrations of Zn, while ‘CDC Impala’, ‘CDC Grimm’, ‘CDC QG-3’ were among the cultivars with the highest concentrations of P (Figure 3).

Higher concentrations of Cu and Mo were observed for specific genotypes in both whole and dehulled seeds (Figure 4). ‘CDC Impala’ and ‘CDC Kermit’ had the highest Cu concentrations in both whole and dehulled seeds, while ‘CDC Grimm’ and the line IBC 1124 had the highest concentrations of Mo in both whole and dehulled seeds (Figure 4). Al and B were essentially only found in the whole seed samples, i.e., in the seed coats, and there were no notable differences among genotypes (Figure 4). Several genotypes had substantially higher concentrations of multiple minerals; this was the case with ‘CDC Impala’, ‘CDC Karim’, ‘CDC Nimble’, ‘CDC Impower’, ‘CDC Grimm’, ‘CDC Imvincible’, ‘CDC Kermit’, ‘CDC QG-3’, ‘CDC SB-4’ and the lines 08M-2 and IBC 1124. Yield was only positively correlated with Fe (r^2^ = 0.66) and was negatively correlated with Cu (r^2^ = 0.64) and B (r^2^ = 0.53) (Figure S2, Table S6 for yield data).

## DISCUSSION

Lentils can contribute significant amounts of minerals to the human diet, and the content will be determined by the cultivar, the form in which it is consumed (whole or dehulled) and, for some minerals, the site where they were grown. To date, the mineral composition of lentils has received little or no attention and has not been a parameter for breeding selection.

With this study, we have established a complete mineral profile of a range of lentil cultivars from the CDC breeding program, thereby giving breeders an idea of the potential for breeding for increasing concentrations of a range of minerals. By analysing samples from diverse growing regions, we were able to determine mineral stability across agro environments. By evaluating both whole seed and dehulled samples, we could determine the potential impact of dehulling on mineral concentrations in raw lentils. We were also able to explore variability across market classes, compared newer material with older cultivars, benchmark against studies conducted elsewhere. In addition to providing information to breeders, these results provide important information for industry partners, as they have expressed interest in the level and stability of mineral profiles, particularly for flavour-potentiating minerals like Na, K and Ca, for marketing purposes.

Few genotypes showed high concentrations of more than one mineral, but in most cases, genotypes were outstanding in one or two. When our found values are compared to previous studies conducted in the last two decades, with some of the CDC cultivars and older varieties cultivated in the same area, range values are about the same (Table S5), implying that breeders have not increased the mineral content of lentils over time. This is not surprising given screening for minerals is time-consuming, destructive, and expensive and not generally a high priority in breeding programs. On a positive note, they have not been unintentionally selected against mineral concentrations leaving room for improvement if desired.

When our study values are compared to the mineral concentration of lentils grown and evaluated in the Mediterranean and Middle East, most ranges are similar. Concentrations of P and Mn were the exception, with some of the reported concentrations in the literature (Table S5) being almost twice the highest values observed in our experiment. In contrast, concentrations of Mn, Fe, and Ca, were about half those observed in our study. The lower mineral concentrations identified in our study, are not necessarily dictated by the genotype entirely, it could be a consequence of the agro environment. Some of the latest and most widely cultivated varieties from the CDC have also exhibited contrasting values of mineral concentrations in a study conducted in Montana (Chen et al., 2022). In that study, ‘CDC Maxim’ had the highest concentration of Fe, Zn and S, while ‘CDC Imvincible’ had the greatest concentration of Se, P and Cu.

Also, Fe was the only mineral with a positive correlation with yield, which implies that increasing yield doesn’t necessarily compromise Fe content.

Introgression from wild related species has been an important approach for improving several characteristics in the lentil crop. Previous research has reported the mineral concentration profile in wild *Lens* species including *Lens lamottei*, *Lens ervoides* and *Lens nigricans* (Gupta et al., 2016, Table S5). Iron concentrations in *Lens lamottei* were slightly superior to our cultivated genotypes. Zn concentration reported from genotypes of these wild species were within the values observed in our analysis. Calcium, Mg and Cu were in higher concentrations compared to the wild species. Although this give us a general idea of the potential of these specific wild genotypes, there are some limitations to these comparisons, given that the Gupta study was conducted in a greenhouse system (Gupta et al., 2016). Genotypes from these and other wild *Lens* species will need to be further study under field conditions to better understand their potential contributions to enhance mineral concentration in lentils.

When compared to raw seed values of other pulses, concentrations of Fe and Se in lentils from our study were similar to those reported for common bean (*Phaseolus vulgaris*), chickpea (*Cicer arietinum*) and pea (*Pisum sativum*) (citations; Table S5). Boron concentrations in lentil were comparable to chickpea. Lentils have higher concentrations of Mg compared to chickpeas.

Similarly, lentil and pea have higher concentrations of Mn compared to chickpea. Chickpea seeds have the highest reported concentrations of Zn, in comparison to those of lentils, common beans and peas, which are similar. Chickpeas also have the highest concentrations of Mg, Ca and K compared to lentils. Lentils have similar concentrations of P compared to chickpeas, and similar concentrations of Cu compared to common beans and chickpeas (Ramirez-Ojeda et al., 2018; Vandemark et al., 2018; Ray et al., 2014).

Some efforts have explored the potential in lentil for Fe and Zn biofortification (Thavarajah, et al. 2009). Genetic variation in seed Fe and Zn concentration and molecular markers related to these traits was observed in a set of 138 cultivated lentil accessions which were grown in four different environments in Saskatchewan (Khazei, et al. 2017).

Some researchers suggest that selection and enhancement of an optimal profile of micronutrients through plant breeding and production site selection may be possible, and practical (Chen et al., 2022; Ray et al., 2014). Other research has found minimal variation among genotypes and large influence of the environment (Vandemark et al., 2018). Results from our study suggest that breeding for an enhanced mineral profile in lentil will be complex as there are no observations of genotypes that are high across the entire set. Mineral biofortification in lentil may be possible, depending on target concentrations. A practical starting point could be with genotypes with high concentrations of more than one mineral, like ‘CDC Impala’, ‘CDC Karim’, ‘CDC Nimble’ (all three red market class), ‘CDC Impower’, ‘CDC Grimm’, ‘CDC Imvincible’, ‘CDC Kermit’ (all four green market class), and ‘SB-4’ (Spanish brown type). If the goal is to enhance concentration values to more than that found in these lines, broader lentil diversity must be explored. In the other hand, potential contributions from wild germplasm could be explored.

What role environment may play in mineral concentrations available to lentil consumers depends on the mineral. A recent field-based study found variation in mineral concentrations across different environments for *most* minerals (Chen et al., 2022). Other studies have shown an impact of the environment on *some* of the minerals they have studied (Ray et al., 2014). And others, have observed no environmental effect, particularly for the Fe concentration of lentils grown in Saskatchewan (Thavarajah et al., 2009). In that study, Fe concentration of raw lentil seeds was not correlated with Fe content in the soil (Thavarajah et al., 2009). In the present study, Na, K, Ca, Mg, Cu, and B appear stable across environments, and thus could be measured with confidence at only one location. This gives plant breeders the possibility to develop germplasm with higher concentrations of this set of minerals that would perform consistently across a range of target environments. In our case, this would be the brown and black soils of Alberta and Saskatchewan, which are generally adequate in mineral composition (Soils of Saskatchewan 2009). In contrast, our study showed that concentrations of Fe, Zn, P, S, Mg and Mn are more affected by the environment and breeding for increased levels of these will require a different, and likely more expensive, strategy than simply testing at a single location. Further studies will also need to be conducted to determine if correlations among minerals concentrations in lentil seeds and specific mineral soil composition of soils in the target growing environment can be established.

Very few studies compare mineral concentrations in whole vs. dehulled seeds but processing of lentil can affect mineral concentrations available to consumers. About 95% of red lentils are eaten dehulled. Hulls account for 6-10% of seed weight but have minerals, among other nutrients, that may only be present in that tissue. Ca, Mg, Mn, B and Fe (for some specific genotypes) are primarily found in the seed coat. When seed coats are removed, these minerals and nutrients are lost to the consumer. In a study conducted in lentil samples from markets in Egypt (Wang et al., 2009), it was found that Ca, Mg, Mn, Cu and Fe were lower in dehulled seeds. The same study showed that K and P concentrations increased with dehulling – i.e. the hull tissue dilutes the mineral content of whole seeds. We found the same result, along with increased concentrations of Ni and Mo in some genotypes. This may have impacts on human nutrition for lentil consumers, as what they receive nutritionally depends on the physical state of the consumed product.

Lentils have antinutritional (chelating) agents that affect the bioavailability of micronutrients, depending on amount in the diet, including phenolic compounds, phytates and oxalic acid (Wang and Daun 2006). Germinating, cooking or fermenting lentils before consumption, can minimize their adverse impact (Margier et al., 2018; Umena et al, 2005). It its known that phytic acid lowers the bioavailability of Fe (Plaza et al., 2021), but Canadian lentils are low in phytic acid (Thavarajah et al., 2011). Dehulling reduces polyphenols and tannins mainly present in the seed coat (Wang et al., 2009) that might have tied up the minerals. For those consuming whole lentils, these compounds are reduced with cooking (Wang et al., 2009) or processing (Ramirez-Ojeda 2018), thus reducing their impact on mineral bioavailability. From analysis of a group of Canadian varieties grown in the past, cooking lentils increased Ca, Cu, and Mn but reduced Fe, K, Mg, P, Zn, and phytic acid (Wang et al., 2009). Al may increase during cooking (Cuciureanu et al., 2000).

Among food products consumed in a daily diet, some may contain minerals classified as heavy metals that are harmful for the organism, including As, Cd, Hg and Pb (Koch et al., 2022). None of these were quantifiable in the samples we analysed. In human diets, we are also exposed to Ba, in drinking water and food (grains, fruits, vegetables, nuts, herbs and spices) the main source of exposure in the Canadian diet (Health Canada, 2012). Barium is a naturally occurring element in the environment, present in Canadian soils, varying widely from 15 to 3000 mg Kg^-1^ (Canadian Council of Ministers of the Environment 2013). About 90-98% is excreted in feces and urine and a tolerable intake of Ba for humans is 0.21 mg of Ba/kg body weight per day (Schroeder et al., 1972). Similarly, Aluminum exposure is inevitable and challenging to quantify or even diagnose human toxicity (Exley 2016). Being Ba and Al present in water, air, soil, and many of the food products we consume, discussing the effects in human health of the amounts of Ba and Al quantified in raw lentils is beyond the limits of this study.

Our work has established a robust mineral profile from a broad range of lentil cultivars and environments. In terms of breeding for enhanced mineral composition in lentils, a biofortification approach could start with the cultivars that exhibited higher concentration values for several of the mineral concentrations evaluated. (example?) This may be a challenging approach considering the ranges in variation observed in this group and the observed wide variation across environments for some minerals. For some specific markets, fortification may be a better approach. Future areas of research could include further exploration of the range of diversity of both cultivated and related wild *Lens* species, or further study of the diverse rooting patterns across the genus *Lens* (Gorim and Vandenberg 2017). By profiling the mineral concentration profiles of segregating interspecific populations with contrasting root phenotypes, we may be able to determine if the rooting profile is associated with mineral uptake into the seed. Further research involving one or more of these approaches may provide a better understanding of whether we are able to enhance the mineral composition of lentil seeds through breeding.

## Acknowledgments

The research was conducted as part of the Application of Genomics to Innovation in the Lentil Economy (AGILE) project [grant: LSP15-8302], a Genome Prairie-managed project funded by Genome Canada, Western Grains Research Foundation, the Province of Saskatchewan, and the Saskatchewan Pulse Growers. The authors are grateful for technical assistance from Barry Goetz, Crop Development Centre, University of Saskatchewan, to analyze micronutrients. Thanks to the pulses team for their technical support.

## Data Availability Statement

The data supporting this study are available online https://knowpulse.usask.ca/experiment/phenomics/Minerals-and-trace-elements-content-of-Canadian-lentil-cultivars.

