## Supplementary material for "Variability in mineral composition of Canadian lentil cultivars": Supplemental Figure S1.docx

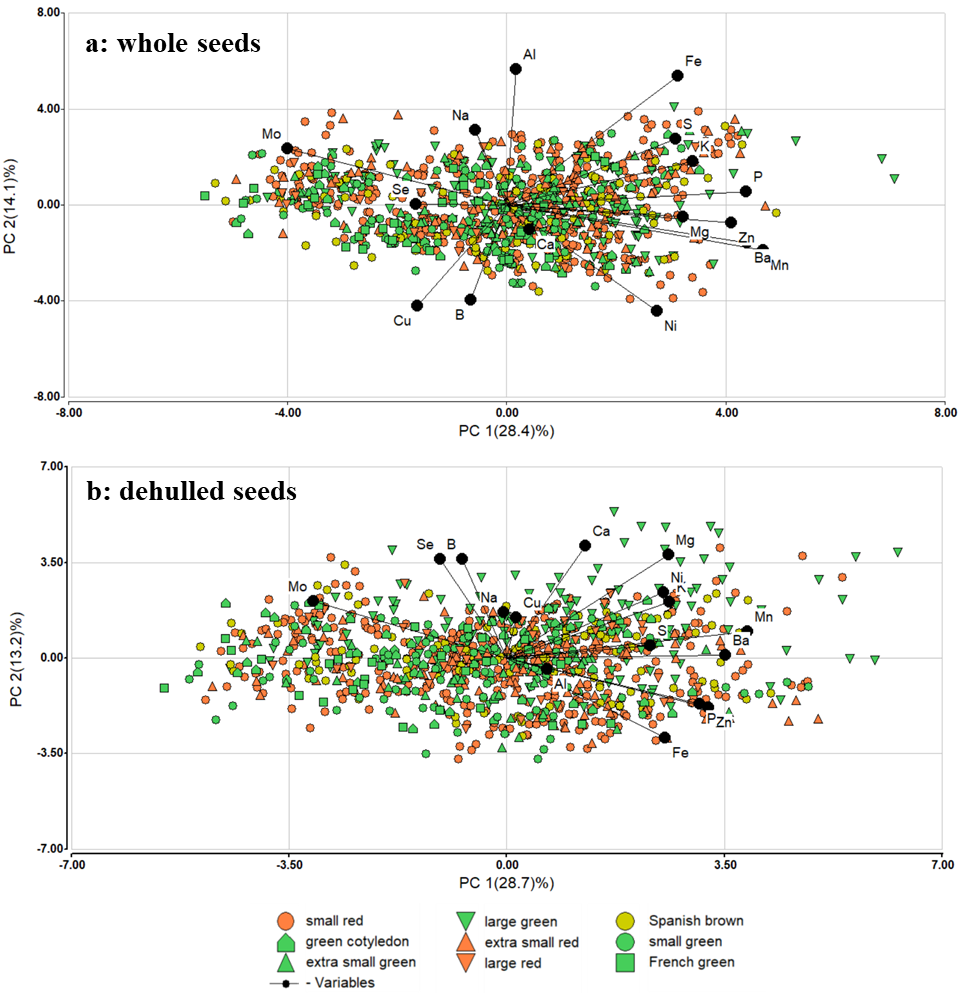

Supplemental Figure S1. Principal component analysis of a: whole seeds and b: dehulled seeds of 34 lentil genotypes according to their market classes grown during 2017-18 in Saskatchewan.
