## Supplementary material for "Variability in mineral composition of Canadian lentil cultivars": Supplemental Figure S2.docx

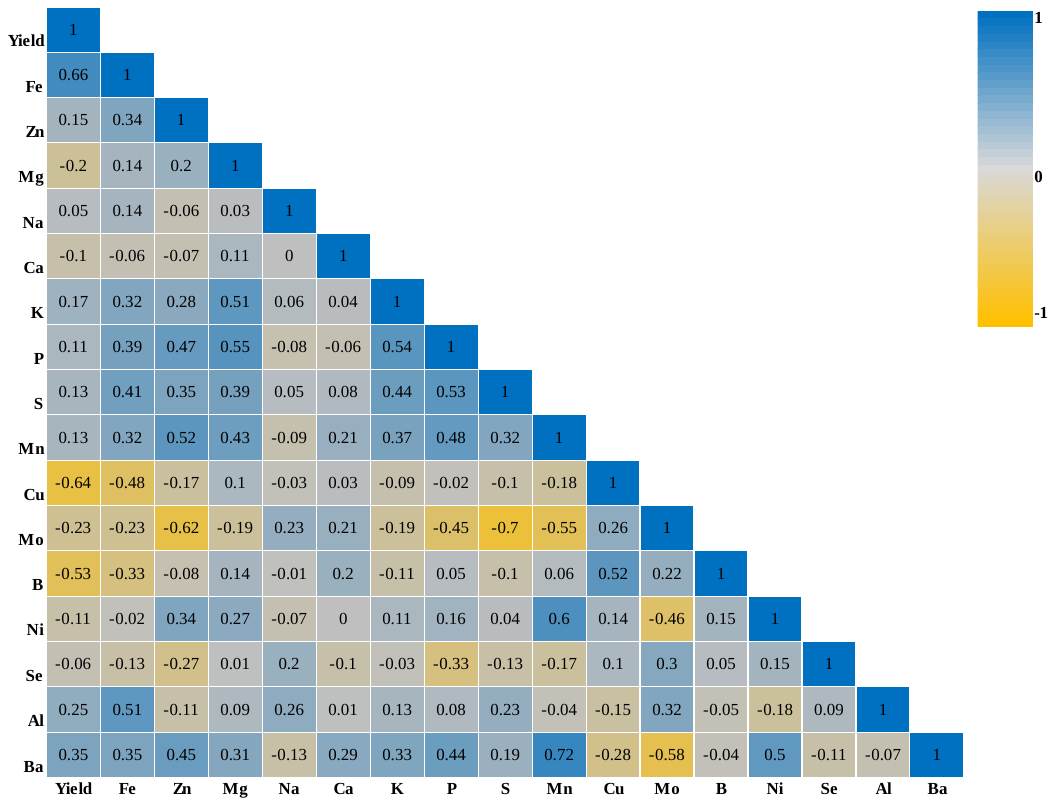

Supplemental Figure S2. Pearson correlations among yield and 16 mineral concentrations evaluated in 34 lentil genotypes grown across 10 site-years in Saskatchewan during 2017-18.
